# Defined human PSC culture conditions robustly maintain human PSC pluripotency through Ca^2+^ signaling

**DOI:** 10.1101/2023.08.08.552440

**Authors:** Ilse Eidhof, Malin Kele, Mansoureh Shasavani, Benjamin Ulfenborg, Dania Winn, Per Uhlén, Anna Falk

**Affiliations:** Department of Neuroscience, Karolinska Institutet, Stockholm, 171 11, Sweden; Department of Medical Biochemistry and Biophysics, Karolinska Institutet, Stockholm, 171 11, Sweden; School of Bioscience, University of Skövde, Skövde, 541 28, Sweden; Department of Experimental Medical Science, Lund University, Lund, 221 84, Sweden

**Keywords:** Human induced pluripotent stem cells, pluripotency, defined PSC culture conditions, Ca^2+^ signaling

## Abstract

Human pluripotent stem cells (hPSCs) have significant potential for disease modeling and cell therapies. However, their clinical applicability is limited due to the need for undefined conditions for PSC cultivation, which increase the risk of immunogenicity, result in batch-to-batch variability and finite scalability. These limitations may be circumvented by xeno-free, defined culture conditions. However, biological processes that preserve robust, homogenous PSCs in defined conditions remain to be characterized. Here, we compared gene expression data from over 100 hPSC cell lines cultivated in undefined and defined culture conditions. Defined culture conditions significantly reduced inter-PSC line variability, highlighting the importance of standardization to minimize PSC biases. This variability is concurrent with decreased germ layer differentiation and increased expression of Ca^2+^-binding proteins. The significance of tightly controlled Ca^2+^ signaling in hPSC pluripotency in defined culture conditions was also confirmed. A deeper understanding of these processes may aid in standardizing defined hPSC culture conditions.

**Graphical abstract:** 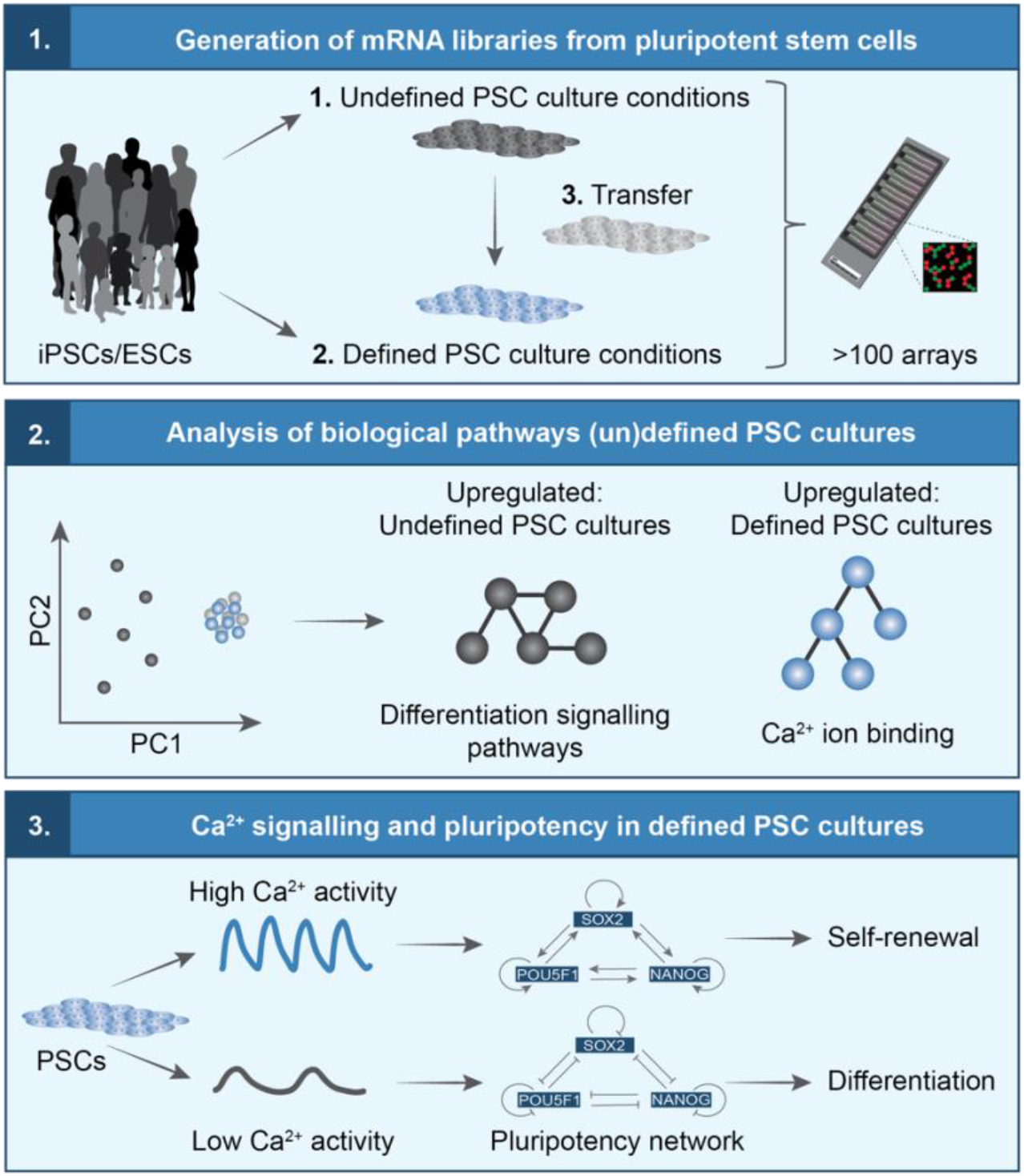

**In Brief:** Eidhof et al. compared gene expression array data obtained from more than 100 hPSC cell lines to study the biological differences between PSCs cultivated under defined and undefined culture conditions.

**Highlights:**

- Defined culture conditions reduce inter-hPSC line variability.
- Defined conditions decrease the expression of germ layer differentiation markers.
- Ca^2+^ signaling associated genes capture specific hPSC states in defined conditions.
- High Ca^2+^ activity drives pluripotency, low Ca^2+^ activity drives differentiation.

## Introduction

The discovery of reprogramming adult skin cells back into induced pluripotent stem cells (iPSCs), which earned the Nobel Prize in 2012, has opened a plethora of possibilities^1^. This remarkable achievement has brought us significantly closer to cell transplantation therapies and gaining deeper insight into the molecular and cellular mechanisms underlying human diseases and early brain development. However, the lack of iPSC standardization, and the difficulties and variability related to maintaining human iPSC pluripotency and self-renewal in undefined culture conditions have restricted the clinical applicability of iPSCs since their discovery. The use of fibroblast feeder cells, Matrigel, and undefined media containing fetal bovine serum (FBS) negatively affect the reproducibility of iPSC lines^2, 3^. They can lead to high batch-to-batch variability in the differentiation potential of iPSC lines between different research laboratories, increase the risk of pathogen contamination and induce immunogenicity^2, 3^. High culture costs associated with undefined iPSC matrices and media further limit their scalability^2, 3^. To address all these challenges, recent efforts have been made by large networks, like the International Society for Stem Cell Research (ISSCR) (https://www.isscr.org/standards-document) and the European College of Neuropsychopharmacology (ECNP) Network to provide researchers with best practices and recommendations for the use of human stem cells in research^4^. Moreover, researchers have explored xeno-free, fully defined iPSC culture matrices, such as laminin 521 (LN-521) and vitronectin, and defined iPSC culture media, such as essential 8 (E8) to facilitate the robust generation and maintenance of iPSCs^2, 3, 5^. Despite these improvements, our knowledge regarding the biological processes that preserve highly robust and homogenous iPSC populations under defined culture conditions is still highly limited. Gaining a deeper understanding of these processes will be crucial in the standardization and improvement of defined iPSC culture conditions. This will ultimately help enhancing the rigor, reproducibility, accurate and unambiguous reporting of iPSC research, and the clinical application of iPSCs in the future.

Here, we performed a comparative analysis of Illumina array data obtained from over 100 human iPSC and embryonic stem cell (ESC) lines cultivated in undefined and/or fully defined conditions. We used this dataset to characterize biological differences between PSCs cultured in (un)defined conditions, and the processes required to maintain PSC pluripotency and self-renewal capacity. Moreover, we shed light on the importance of standardization to minimize biases and variations other than those caused by differences in the genetic background between PSC lines.

## Results

### Defined culture conditions promote greater uniformity among PSC lines

To investigate pathways involved in maintaining PSC pluripotency and self-renewal in defined culture conditions, we utilized our substantial collection of Illumina gene expression array data obtained from over 100 healthy human control iPSC, neurogenetic patient iPSC and ESC lines. These PSC lines were either cultivated and maintained under undefined or fully-defined culture conditions, or transferred from undefined to defined culture conditions (**Fig. 1A-B**). All PSC lines were cultured in the presence of bFGF. The iPSCs showed a morphology similar to that of human ESCs, with large nuclei, compact cytoplasm, and sharp edges. Moreover, all iPSCs/ESCs included in this study passed the PluriTest criteria for pluripotency, as evidenced by their Pluripotency and Novelty scores, expression of PSC markers, and normal karyotypes (**Table S1**). A comprehensive overview of the samples and culture conditions of the three different groups is presented in **Fig. 1B**, whereas a complete list of all PSC lines and samples is provided in **Table S1**.

**Figure 1.**
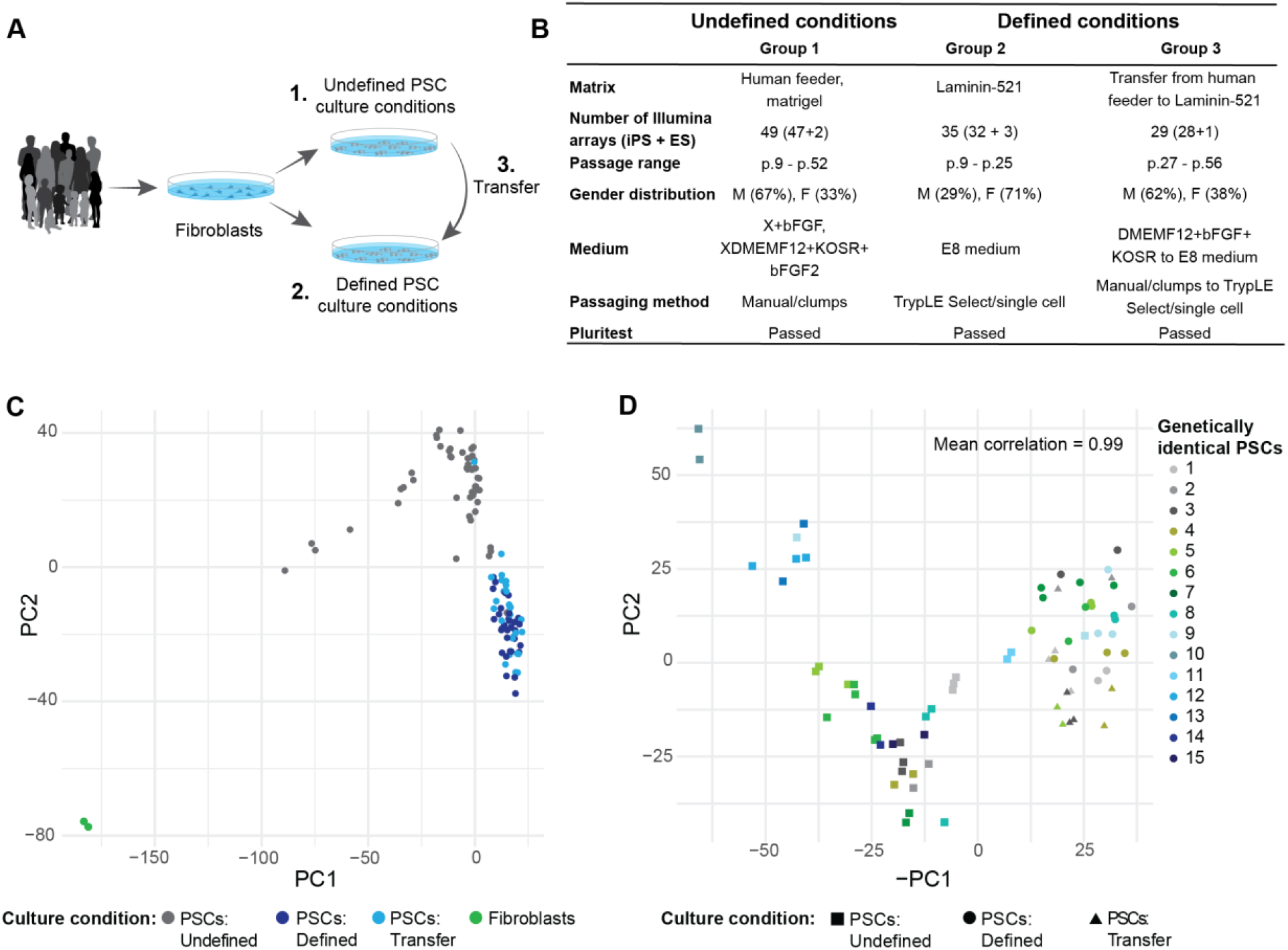
Defined culture conditions promote greater uniformity among PSC lines compared to undefined culture conditions. **A.** Schematic representation of the three groups included in the analysis. **B.** Table summarizing the exact culture conditions and number of PSC lines for each group. **C.** PCA visualization of PSC lines derived and cultured in 1. Undefined culture conditions (in grey), 2. Defined culture conditions (in dark blue), 3. Transferred from undefined to defined culture conditions (in light blue) and 4. Fibroblasts (in green). **D.** PCA visualization of PSC lines derived and cultured in 1. Undefined culture conditions (in squares), 2. Defined culture conditions (in circles) and 3. Transferred from undefined to defined culture conditions (in triangles). The color codes represent different technical or biological replicates. When there is a similar color code in one of the three groups (undefined, defined or transfer), the data originates from either different iPSC clones of the same individual, iPSCs from monozygotic twins, or technical replicates of the same PSC line.

We performed principal component analysis (PCA) to identify the factors contributing to variability between PSC line samples in the Illumina array dataset. We also included Illumina array data from two human skin fibroblast cell lines. Our analysis indicated that the largest variability among the four different groups (Principal Component 1 (PC1), 20%), was determined by cell type (**Fig. 1C**). PSCs formed a separate cluster away from the fibroblast cell lines, confirming that PSC lines of different origin were similar to each other. PC1 also displayed partial separation between PSCs cultured under undefined conditions (**Fig. 1C**, in grey). PC2, which accounted for 13% variability in the dataset, separated a more widespread, undefined PSC culture condition group (in grey) from a more homogenous, fully defined PSC culture condition group (dark blue) (**Fig. 1C**). Importantly, PSCs that were transferred from undefined to fully defined culture conditions (light blue) demonstrated reduced variability within PSC lines compared to their undefined counterparts, and largely overlapped with PSCs cultures under defined conditions (**Fig. 1C**).

Next, we focused on the PSC lines in PCA by excluding the fibroblast samples from the analysis. Our analysis revealed that the primary source of variability explained by PC1 (20%) could be attributed to transcriptional differences between PSCs cultivated under defined and undefined conditions (**Fig. S1A**). Other potential confounding variables in the dataset, such as the diagnosis, sex of the donor, passage number, and/or origin of the PSC line, did not contribute significantly to the observed variability in the PCA (**Fig. S1B-E**). Consistently, we observed that both PSCs cultured in defined conditions (dark blue) or those transferred to defined culture conditions (light blue) exhibited higher homogeneity, and formed tightly clustered groups compared to their undefined counterparts (gray) (**Fig. S1A**). Thus, despite meeting the quality criteria for pluripotency, the PSCs grown under undefined culture conditions displayed considerably larger variability compared to those cultured in defined conditions.

To further investigate the greater heterogeneity observed in samples from undefined culture conditions, we examined whether genetically identical samples exhibited closer clustering and higher correlation in defined culture conditions compared to undefined conditions (**Fig. 1D**). These genetically identical PSC lines included iPSCs derived from monozygotic twins, iPSC clones originating from the same donor, and/or technical duplicates from the same PSC line. Notably, none of the genetically identical PCS sample duplicates were shared between the defined and undefined PSC culture conditions. Genetically identical PSC lines clustered tightly together in PCA, with a mean correlation of 0.99 both for PSCs cultured under defined and undefined conditions (**Fig. 1D**). This suggests that genetically identical PSC samples are largely homogenous regardless of their exact culture conditions. Thus, the greater heterogeneity observed among PSCs under undefined culture conditions can be attributed to a higher overall inter-PSC line variability.

In conclusion, defined culture conditions promoted homogeneity between PSC lines.

### Pluripotency is maintained in defined culture conditions via inhibition of signaling pathways that regulate germ-layer differentiation

We next performed analysis of differentially expressed genes (DEGs) to determine the biological processes and pathways that contribute to increased inter-PSC line homogeneity under defined culture conditions. A total of 313 DEGs (with a fold change greater than 1 and p < 0.05) were identified between the undefined and defined PSC culture condition groups (**Fig. 2A**), consisting of 172 downregulated and 141 upregulated genes (**Fig. S2A-B**). A total of 345 DEGs were identified between PSCs cultured under undefined conditions and those transferred from undefined to defined culture conditions (**Fig. 2A**), including 221 downregulated and 124 upregulated genes (**Fig. S2A-B**). Interestingly, only 3 genes were found to be differentially expressed exclusively between PSCs cultured in defined conditions and PSCs transferred from undefined to defined culture conditions (**Fig. 2A**), including two downregulated and one upregulated gene(s) (**Fig. S2A-B**). This indicates a high degree of similarity between these two groups. Indeed, a high proportion of DEGs (227 of the 313 DEGs, 72.5%) overlapped between PSCs under defined culture conditions and those transferred from undefined to defined culture conditions (**Fig. 2A**). Among these overlapping DEGs, 146 and 81 were up- and down-regulated, respectively (**Fig. S2A-B**).

**Figure 2.**
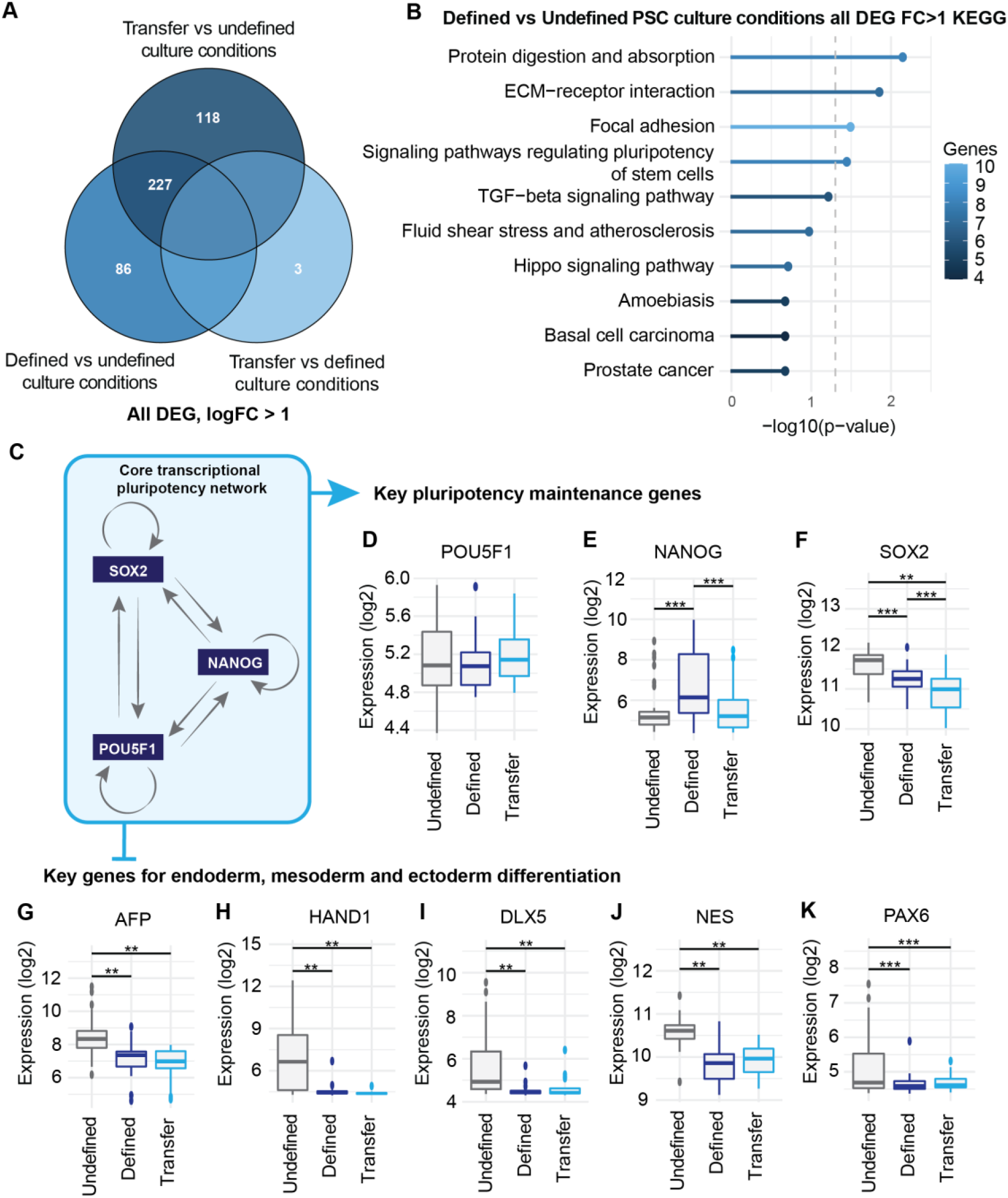
Pluripotency is maintained in defined culture conditions via inhibition of signaling pathways that regulate germ-layer differentiation. **A.** VENN diagram showing the total number of DEGs passing a log foldchange > 1 between the indicated PSC culture conditions. **B.** KEGG pathway analysis on DEGs between PSC established in defined and undefined culture conditions. **C.** Schematic overview of the core transcriptional network regulating stem cell pluripotency. **D-F.** Log2 microarray expression values of *POU5F1* (D), *NANOG* (E) and *SOX2* (F) that make up the core network in C. for the indicated PSC culture conditions. **G-K.** Log2 microarray expression values of the endodermal marker *AFP* (**G**), mesodermal markers *HAND1* (**H**) and *DLX5* (**I**), and ectodermal markers *NES* (**J**) and *PAX6* (**K**) for the indicated PSC culture conditions. **D-K:** Statical testing: one-way ANOVA followed by Tukey HSD for post-hoc testing: **p < 0.01, ***p < 0.001.

We continued asking whether the 313 DEGs between undefined and defined PSC culture conditions were significantly enriched for biological pathways in the Kyoto Encyclopedia of Genes and Genomes (KEGG) database. The DEGs showed among others significant enrichment in the KEGG annotations: protein digestion and absorption (p < 0.01), ECM-receptor interaction (p < 0.05), focal adhesion (p < 0.05), and signaling pathways regulating the pluripotency of stem cells (p < 0.05) (**Fig. 2B**). A similar analysis was conducted for 345 DEGs between the transfer and undefined PSC culture groups, where similar processes were identified: focal adhesion (p < 0.01), signaling pathways regulating the pluripotency of stem cells (p < 0.01), p53 signaling pathway (p < 0.05), and protein digestion and absorption (p < 0.05) (**Fig S2C**).

These findings prompted us to delve deeper into the genes that constitute the core transcriptional network responsible for maintaining pluripotency in the Illumina array dataset (**Fig. 2C**). Notably, no significant changes in *POUF51* expression were observed (**Fig. 2D**). However, *NANOG* was significantly upregulated (p < 0.001) under the defined culture conditions (**Fig. 2E**). *SOX2* was significantly downregulated in both PSCs cultured in defined conditions (p < 0.001) and in PSCs transferred from undefined to defined culture conditions (p < 0.01) (**Fig. 2F**). These findings suggest that the PSC culture conditions affect the expression of genes that constitute the pluripotency network.

To examine the downstream effects of DEGs in the pluripotency network, we conducted an analysis of the expression of early mesodermal, endodermal, and ectodermal fate genes commonly used in embryoid body assays. Interestingly, a significant proportion of these genes, including the endodermal marker *AFP* (p < 0.01), mesodermal markers *HAND1* (p < 0.01) and *DLX5* (p < 0.01), and ectodermal markers *NES* (p < 0.01) and *PAX6* (p < 0.001), were significantly downregulated in PSCs cultured in defined conditions and in PSCs transferred from undefined to defined culture conditions (**Fig. 2G-K**). No significant changes in the expression of the endodermal markers *AMY1A*, *FOXA2*, and *PDX1*; mesodermal marker *FLT1*; and ectodermal markers *GFAP* and *SOX1* were observed (**Fig. S2D-I**).

These findings indicate that the defined culture conditions facilitate the maintenance of a homogeneous population of PSCs. The larger variability in PSCs maintained under undefined culture conditions may be attributed to the presence of a heterogeneous population consisting of both PSCs and PSCs that have initiated or committed to differentiation. Importantly, despite this variability, all PSCs passed the pluripotency quality criteria. This suggest that conventional methods used to assess pluripotency are limited in sensitivity, since PSC cultures with pluripotent, committed and partly differentiated cells passes as pluripotent.

### DEGs identified between defined and undefined PSC culture conditions are strongly interconnected at the protein level and involved in shared biological processes

The understanding of protein-protein interactions, both in terms of their physical presence and functional relationships, is crucial for elucidating the mechanisms underlying the maintenance of robust, homogenous PSC populations under defined culture conditions. To evaluate whether our list of DEGs in the defined culture conditions showed significant molecular connectivity, we first collected protein-protein interactions (PPI) using the Genemania plugin in Cytoscape^6^. Then, we explored which PPI interactions were more enriched among our list of DEGs compared to what would be expected by chance alone through a bootstrap analysis^7^. These significant PPI interactions were then integrated into a reference network containing 247 nodes and 645 edges (**Fig. 3**) representing various types of protein interactions: physical protein interactions (p < 0.0001), predicted protein interactions (p < 0.0001), shared protein domains (p < 0.0001), pathway interactions (p < 0.0001), and co-localized proteins (p < 0.0001) (**Fig. S3A**). The resulting PPI network included a large module comprising 172 nodes and 618 edges, and six smaller modules comprising two or more nodes. A total of 59 proteins were not associated with any of the modules (**Fig. 3B**). The physical interaction enrichment score^7^ was recalculated for the complete integrated PPI network. Proteins comprising this network showed significant molecular connectivity (p < 0.0001), with an interaction enrichment score of 1.7 (**Fig. 3A**).

**Figure 3.**
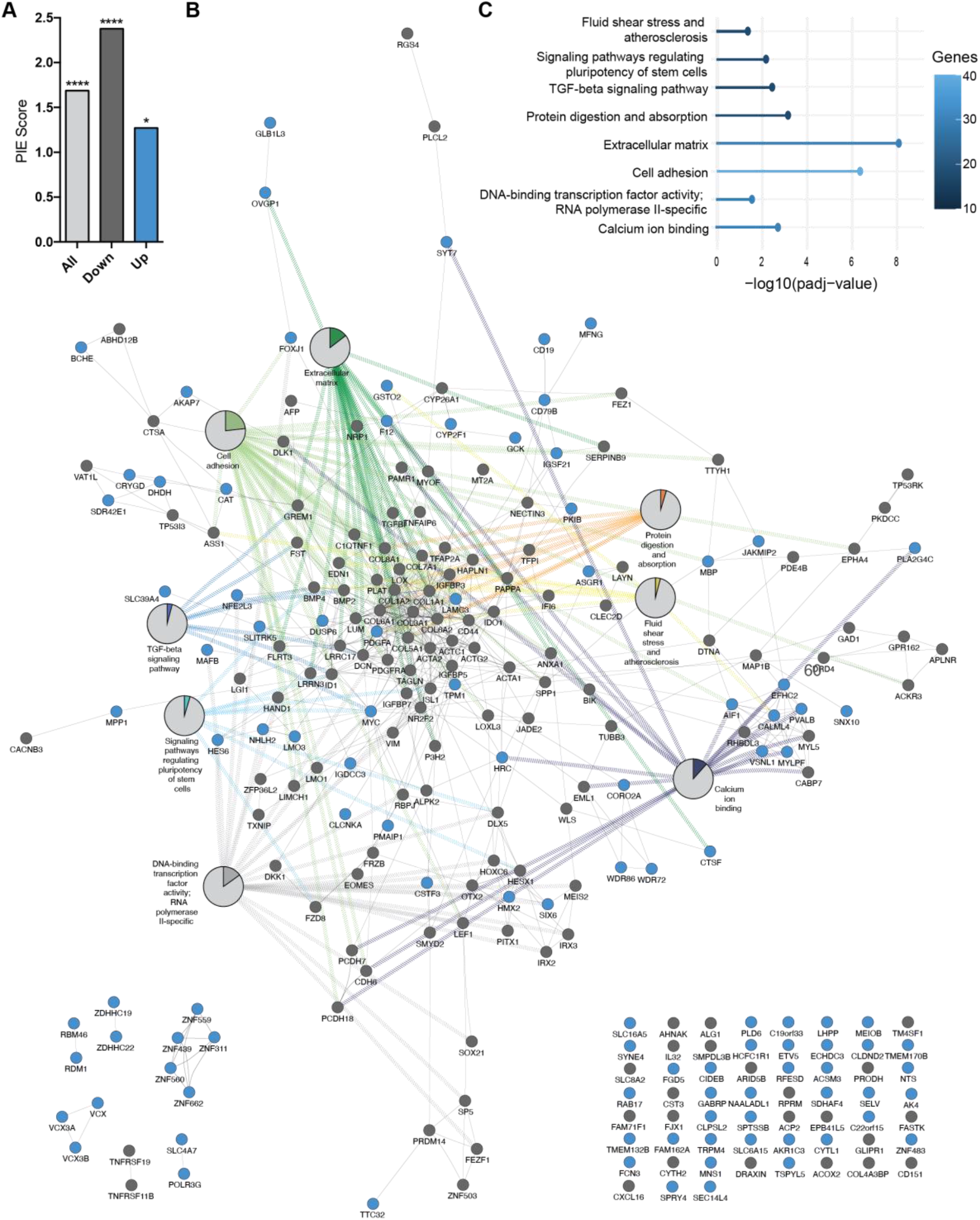
DEGs between defined and undefined PSC culture conditions are strongly interconnected at protein level and function in shared biological processes. **A.** Physical interaction enrichment (PIE) score of all (in light grey), downregulated (in dark grey) and upregulated (in blue) DEGs between defined and undefined PSC culture conditions. *p < 0.05, ****p < 0.0001, based on 10,000 repetitions. **B.** Interaction network of proteins differentially expressed between defined and undefined PSC culture conditions and their significant associated GO terms. Pie charts display the percentage of proteins within the PPI network associated with the indicated GO term. **C.** Significantly enriched GO terms for the large PPI network displayed in B.

We next examined whether the identified PPI modules represent specific biological processes using Gene-Ontology (GO) and KEGG pathway analyses. The large protein module exhibited significant enrichment for several processes, including signaling pathways regulating the pluripotency of stem cells (p < 0.01), TGF-beta signaling (p < 0.01), cell adhesion (p < 0.0001), extracellular matrix (p < 0.0001), DNA-binding transcription factor activity: RNA polymerase II-specific (p < 0.05), protein digestion and absorption (p < 0.001), fluid shear stress and atherosclerosis (p < 0.05) and calcium ion (Ca^2+^)-binding (p < 0.01). Interestingly, both upregulated (blue) and downregulated (dark gray) DEGs were jointly represented within this network, indicating a biological overlap (**Fig. 3B**). Nevertheless, when the PIE scores of upregulated and downregulated DEGs in defined culture conditions were calculated separately, the PIE score of downregulated genes markedly increased to 2.4 (p < 0.0001), whereas that of upregulated genes decreased to 1.3 (p < 0.05), compared to all DEGs (**Fig. 3A**). This trend was also observed in the PIE score analysis of distinct PPI categories (**Fig. S3B-C**). The results revealed that physical protein interactions (p < 0.0001), predicted protein interactions (p < 0.0001), shared protein domains (p < 0.0001), pathway interactions (p < 0.01), and colocalized proteins (p < 0.0001) were significantly enriched among the downregulated genes (**Fig. S3B**). However, none of the unique PPI categories was significantly enriched among the upregulated genes under defined PSC culture conditions (**Fig. S3C**).

We further investigated whether different biological processes at the protein network level were specifically associated with down- and upregulated genes under the defined PSC culture conditions. For this purpose, separate PPI networks were built for the down- and upregulated genes (**Fig. 4A, C**).

**Figure 4.**
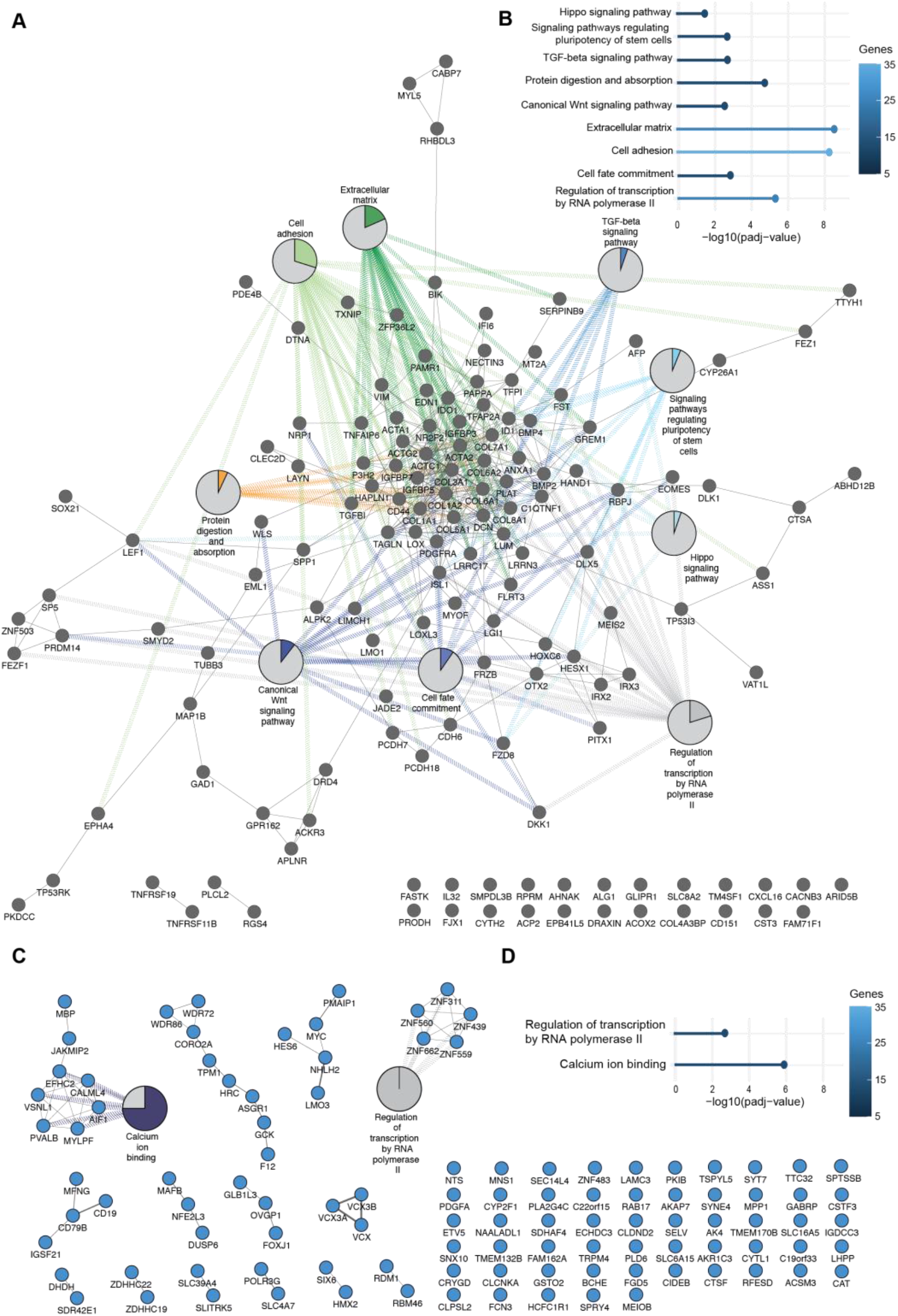
Up- and downregulated genes between defined and undefined PSC culture conditions are strongly interconnected at protein level but are associated with distinct biological processes. **A-B:** Analysis performed with downregulated genes in defined PSC culture conditions. **A.** Interaction network of proteins downregulated in defined PSC culture conditions and their significant associated GO terms. Pie charts display the percentage of proteins within the PPI network associated with the indicated GO term. **B.** Significantly enriched GO terms for the large PPI network displayed in A. **C-D:** Analysis performed with upregulated genes in defined PSC culture conditions. **C.** Interaction network of proteins upregulated in defined PSC culture conditions and their significant associated GO terms. Pie charts display the percentage of proteins within the PPI network associated with the indicated GO term. **D.** Significantly enriched GO terms for the PPI network displayed in C.

The downregulated genes under the defined PSC culture conditions formed a large and strongly interconnected PPI network along with two small PPI modules, comprising 141 nodes and 414 edges (**Fig. 4A**). Additionally, 23 nodes did not interact with any of the modules in the PPI network (**Fig. 4A**). GO and KEGG pathway analyses of the large, downregulated PPI module revealed a striking overlap with significantly associated GO terms linked to the PPI network composed of all DEGs in **Fig. 3B**. Signaling pathways regulating the pluripotency of stem cells (p.adj < 0.01), TGF-beta signaling (p.adj < 0.01), protein digestion and absorption (p.adj < 0.0001), extracellular matrix (p.adj < 0.0001), cell adhesion (p.adj < 0.0001), and negative regulation of transcription by RNA polymerase II (p.adj < 0.0001) were all significantly associated with this PPI module (**Fig. 4A-B**). Furthermore, the PPI module containing the majority of downregulated genes was also significantly enriched with unique KEGG pathway terms linked to pluripotency and differentiation, such as the Hippo signaling pathway (p.adj < 0.05), canonical Wnt signaling pathway (p.adj < 0.01), and cell fate commitment (p.adj < 0.001) (**Fig. 4A-B**).

The upregulated genes under the defined PSC culture conditions formed 14 small PPI modules consisting of 106 nodes and 67 edges (**Fig. 4C**). Additionally, 55 nodes did not interact with any of the fourteen modules in the PPI network (**Fig. 4C**). GO and KEGG pathway analyses were performed for modules consisting of five or more nodes. Similar to the downregulated genes, the significantly associated GO terms overlapped with those associated with the PPI network consisting of all DEGs in **Fig. 3B**. One module consisting of five proteins was significantly enriched in transcription regulation by RNA polymerase II (p.adj < 0.01) (**Fig. 4C-D**). However, this term was also enriched in the PPI network comprising downregulated DEGs in **Fig. 4A**. Another PPI module comprising eight upregulated proteins showed significant enrichment for Ca^2+^-binding (p.adj < 0.0001). Interestingly, although we observed downregulated Ca^2+^-binding proteins in the PPI network of all DEGs in **Fig. 3B**, this term was not significantly enriched in the PPI network consisting of downregulated proteins. This suggests that genes associated with Ca^2+^-binding are uniquely enriched among the proteins upregulated under defined PSC culture conditions, and may have a role in the maintenance of pluripotency.

Taken together, these findings demonstrate that differentially-expressed proteins under defined PSC culture conditions are involved in shared biological networks and processes. Furthermore, specific processes are uniquely enriched among either downregulated or upregulated proteins, highlighting their potential significance in cellular maintenance of pluripotency.

### Ca^2+^ activity and Ca^2+^-binding proteins that are upregulated in defined PSC culture conditions are linked to the pluripotent state

Gene expression and PPI network analyses clearly indicated that downregulated genes under the defined PSC culture conditions were associated with PSC differentiation. Therefore, we sought to explore the potential connection between genes associated with Ca^2+^-binding, uniquely enriched at the protein level among the upregulated genes in defined PSC culture conditions, and the pluripotent state. To test this hypothesis, iPSCs were differentiated under defined PSC culture conditions. In parallel, iPSCs obtained from healthy individuals (Ctrl5^8^, Ctrl7^9, 10^, Ctrl9^5^, Ctrl10^5^ and Ctrl14^11^) were cultured on Laminin-521 in defined E8™ and E6™ PSC media. Compared to E8™ medium, E6™ medium lacks two important growth factors, TGFβ and bFGF, which are required for maintaining pluripotency. After seven days of culture, E6 iPSCs showed clear signs of differentiation compared with their E8 counterparts. E6 iPSCs were heterogeneous, showing either regular iPSC morphology, larger, more flattened, and/or radially organized and more elongated morphology compared to iPSCs cultured in E8 medium (**Fig. S4A-B**). Moreover, iPSCs cultured in E6 proliferated slower than those cultured in E8 medium. Upon harvesting on day 7, the total number of cells in the E6 medium was considerably lower than that in the E8 medium, despite equal seeding numbers and densities (**Fig. S4C**).

Since iPSCs cultured in E6 medium for 7 days displayed typical signs of differentiation, we employed flow cytometry to assess the expression of proteins constituting the core pluripotency regulatory network. The iPSCs cultured in E6 medium showed heterogeneous expression of POU5F1 and NANOG compared to those cultured in E8 medium (**Fig. S4D**). E6 iPSCs showed an increased population of cells with low or no expression of POU5F1 and NANOG, whereas SOX2 expression was slightly increased in E6 compared to that in E8 culture conditions (**Fig. S4D**). These results indicate that the pluripotency regulatory network was affected following 7 days of iPSC culture in E6 medium.

To investigate the association between the effects on the pluripotency network and the upregulation of genes involved in early differentiation, we conducted RT-qPCR analysis to measure the expression of *AFP*, *HAND1*, *DLX5*, *NES*, and *PAX6*. *AFP* was excluded from further analysis since its expression levels were too low to be reliably detected by RT-qPCR. The mesodermal markers *HAND1* and *DLX5* were significantly and consistently upregulated in all five E6 iPSCs cell lines tested (**Fig. S4E**). The ectodermal marker *NES* was upregulated in E6 compared to E8 culture conditions, although it did not pass the correction for multiple testing (**Fig. S4E**). Variable results were observed for the ectodermal marker *PAX6*, with some iPSC lines showing increased and others decreased *PAX6* expression when cultured in E6 compared to E8 media (**Fig. S4E**). These findings suggest that 7 days of iPSC culture in E6 medium pushed iPSCs towards a state of unguided differentiation.

Following the establishment of a defined culture condition that mimics the undefined condition by allowing for a heterogenous population of undifferentiated and differentiated iPSCs, we investigated whether the genes that form the Ca^2+^-binding protein module in **Fig. 3B** are differentially expressed upon iPSC differentiation. A simplified representation of this module is shown in **Fig. 5A**. We focused on a subset of the module that showed high interconnection between nine Ca^2+^ ion-binding genes at the protein level, sharing similar protein domains. Using this module, we tested the expression after 7 days of culture in E6 and E8 media of four genes associated with Ca^2+^-binding (*CALML4*, *AIF1*, *PVALB* and *VSNL1*) that were upregulated, and three (*RHBDL3*, *MYL5* and *CABP7*) that were downregulated under defined PSC culture conditions. Interestingly, the expression patterns of all tested genes confirmed the Illumina array data. Thus, all genes that were upregulated under the defined culture conditions were significantly downregulated after 7 days of culture in E6 media compared to those in E8 media (**Fig. 5B**). In contrast, all genes that were downregulated under the defined culture conditions were significantly upregulated after 7 days of culture in E6 media compared to E8 media (**Fig. 5C**). These results indicate that Ca^2+^-binding genes upregulated under the defined PSC culture conditions are associated with the pluripotent state, whereas those downregulated are linked to PSC differentiation.

**Figure 5:**
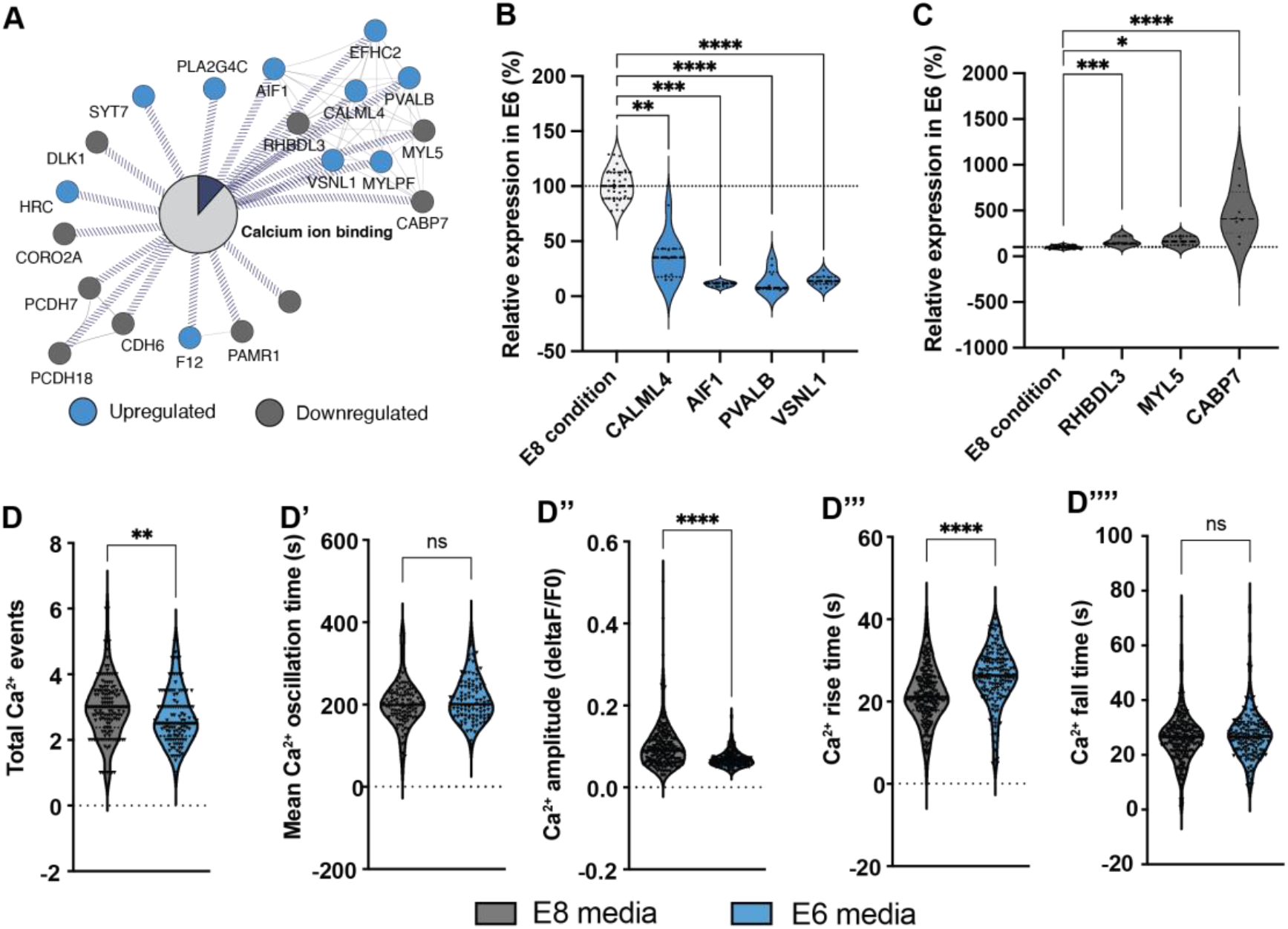
Ca^2+^ activity and Ca^2+^-binding genes that are upregulated in defined iPSC culture conditions are linked to the pluripotent state. **A.** Isolated Ca^2+^-binding protein interaction network based on Fig. 3B. Blue: upregulated proteins in defined iPSC culture conditions. Grey: downregulated proteins in defined iPSC culture conditions. **B.** Upregulated Ca^2+^-binding genes in defined culture conditions, are downregulated when iPSCs start leaving the pluripotent state. QPCR gene expression data is normalized to E8 condition (white). **C.** Downregulated Ca^2+^-binding genes in defined culture conditions, are upregulated when iPSCs start leaving the pluripotent state. QPCR gene expression data is normalized to E8 condition (white). **D.** Live Ca^2+^ imaging of iPSCs cultured in E8 media (in grey) or E6 media (in blue) in baseline conditions measuring the total number of Ca^2+^ transients (D), mean oscillation period (D’), Ca^2+^ peak amplitude (D’’, change of fluorescence over baseline fluorescence), Ca^2+^ peak rise time (D’’’) and Ca^2+^ peak fall time (D’’’’). Pooled data of 5 different iPSC control lines plotted in volcano plots, line indicate median. *p < 0.05, **p < 0.01, ***p < 0.001, ****p < 0.0001, ns: not significant.

Since the expression and function of specific Ca^2+^-binding proteins may depend on the overall spontaneous Ca^2+^ activity in a cell, we investigated whether iPSCs cultured in E6 medium displayed changes in their baseline Ca^2+^ activity compared to iPSCs cultured in E8 medium. Upon cultivation in E6 media, iPSCs showed significant reductions in the number of spontaneous Ca^2+^ transients (p < 0.01) and the Ca^2+^ peak amplitude (p < 0.0001) and a significant increase in the Ca^2+^ peak rise time (p < 0.0001) (**Fig. 5D, 5D’’, 5D’’’**). However, other parameters, such as the mean Ca^2+^ oscillation period and Ca^2+^ peak fall time, were not significantly affected (**Fig. 5D’, 5D’’’’**). This suggests that the spontaneous Ca^2+^ activity is hampered when iPSCs begin to differentiate.

### Ca^2+^ activity is regulated differently in iPSCs that leave the pluripotent state

The spontaneous Ca^2+^ activity can be controlled by a tight interaction between ion channels, adhesion molecules, and receptor proteins^12^.

To assess the impact of unguided iPSC differentiation on voltage-gated Na^2+^ and Ca^2+^ ion channels, iPSCs cultured in E6 and E8 media were depolarized, exposing cells to a high extracellular concentration of potassium chloride (KCl). Approximately 30% of iPSCs cultured in E8 medium responded to the high KCl stimuli (**Fig. 6A, in grey**). In contrast, only 9% of iPSCs cultured in E6 medium responded (**Fig. 6A, in blue**). iPSCs cultured in E6 media that responded to KCl stimuli had a significantly reduced Ca^2+^ peak amplitude (p < 0.0001) compared to that of iPSCs cultured in E8 media (**Fig. 6A’-A’’**). These results indicate that the cellular components involved in depolarization, such as voltage-gated Na^2+^ and Ca^2+^ channels, become less important in the regulation of Ca^2+^ activity during unguided iPSC differentiation.

**Figure 6:**
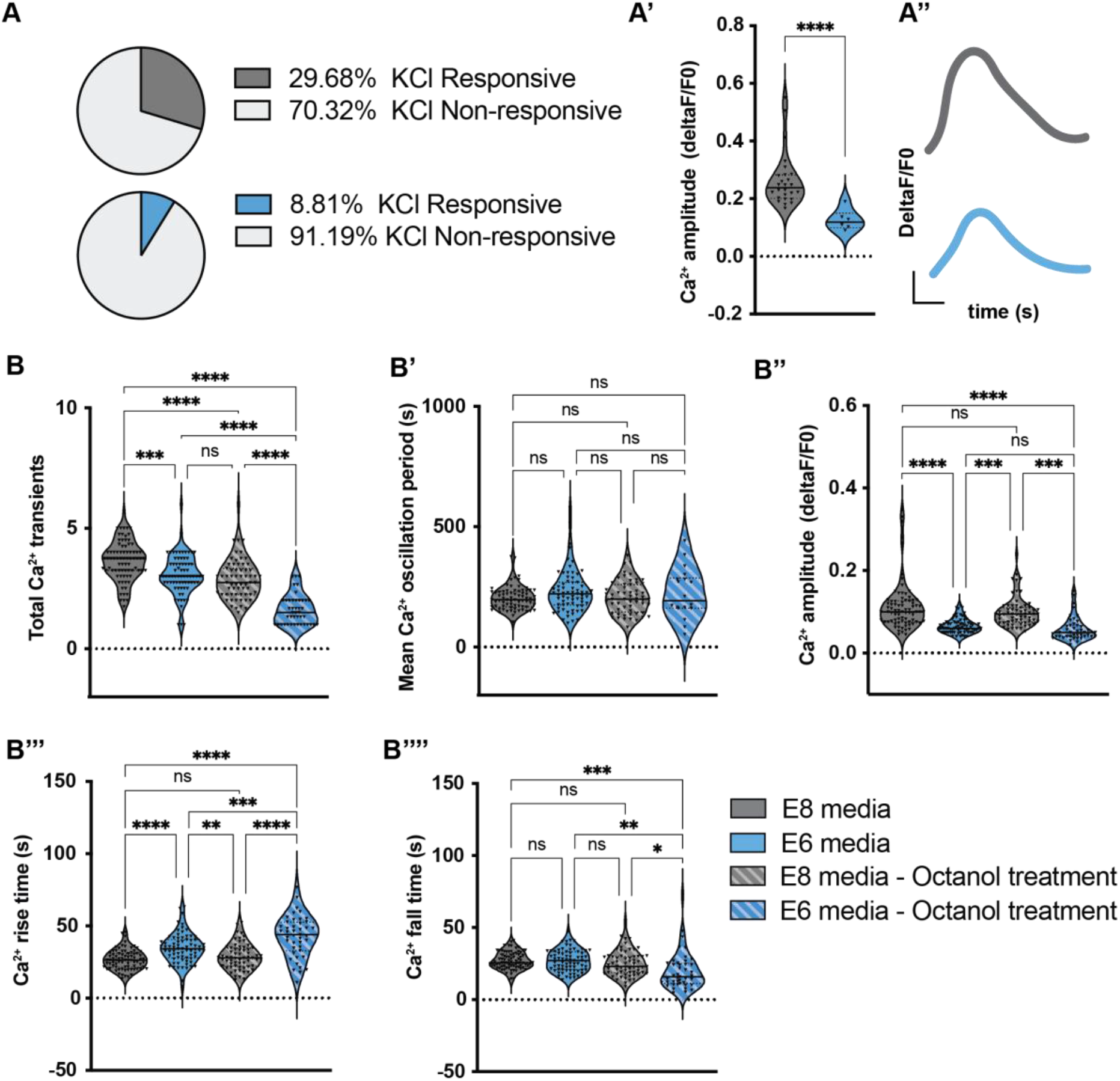
Ca^2+^ activity is regulated differently in iPSCs that leave the pluripotent state. **A.** Pie chart displaying the percentage of cells cultured in E8 (in grey) and E6 (in blue) media that respond to depolarization stimuli. **A’** Ca^2+^ peak amplitude (change of fluorescence over baseline fluorescence) in response to depolarization stimuli. **A’’** Representative Ca^2+^ peak of iPSCs cultured in E8 media (in grey) and E6 media (in blue) in response to depolarization stimuli. **B.** Live Ca^2+^ imaging of iPSCs cultured in E8 media (in grey) or E6 media (in blue) in baseline conditions and upon octanol treatment (diagonal lines) measuring the total number of Ca^2+^ transients (B), mean Ca^2+^ oscillation period (B’), Ca^2+^ peak amplitude (B’’, change of fluorescence over baseline fluorescence), Ca^2+^ peak rise time (B’’’) and Ca^2+^ peak fall time (B’’’’).

We next investigated the influence of cell-cell contact, such as gap junctions, on Ca^2+^ activity of iPSCs and that of differentiating iPSCs. IPSCs cultured in both E8 and E6 media displayed a significant reduction in spontaneous Ca^2+^ activity upon treatment with the gap junction blocker, octanol-1 (**Fig. 6B**). This effect was more pronounced for iPSCs cultured in E6 medium, which showed a significant reduction in the total number of Ca^2+^ transients (p < 0.0001), a significant increase in the Ca^2+^ peak rise time (p < 0.001), and a significant decrease in the overall Ca^2+^ peak fall time (p < 0.01) after octanol-1 treatment (**Fig. 6B, B’’’ and B’’’’**). This demonstrated that cell-cell interaction is required to regulate Ca^2+^ activity at the iPSC stage and during early iPSC differentiation. However, the specific types and number of intercellular contacts influencing Ca^2+^ activity may change as iPSCs differentiate.

## Discussion

In this study, we have performed a comparative analysis of Illumina array data obtained from over 100 healthy human individuals PSC lines derived and cultured in undefined and/or fully-defined PSC culture conditions to elucidate the intracellular differences between pluripotent stem cells cultivated under defined and undefined PSC culture conditions.

### Defined hPSC culture conditions reduce variability observed between different PSC lines

We showed that PSCs could be generated under both defined and undefined culture conditions, as evidenced by all PSC lines passing standardized pluripotency quality controls. Deriving PSC lines under defined conditions, however, largely reduces inter-line PSC variability, regardless of the associated sex, origin of the cell line, passage number, or disease state of the donor. The high level of heterogeneity between PSCs under undefined culture conditions may significantly affect their reproducibility and standardization of their differential potential compared with PSCs cultured under defined conditions.

Our findings are consistent with those reported in earlier independent, smaller-scale studies that have also demonstrated benefits of defined culture conditions. Accordingly, hESCs cultured on a defined recombinant LN-521 matrix exhibited less differentiation, improved cell-matrix adhesion, and increased proliferation compared to other matrices such as Matrigel^3, 13–15^. LN-521 was suggested to enhance stabilization of the expression of pluripotency genes, and support PSC proliferation and self-renewal during long-term culture/passaging^3, 13–15^.

The reduced variability observed between our PSC lines under defined culture conditions coincided with the reduced expression of germ layer differentiation markers, and increased expression of the pluripotency marker *NANOG*. Previous studies have demonstrated a direct effect of TGFβ signaling on the regulation and expression of *NANOG*^16^. In contrast to undefined PSC media, the defined PSC culture media used in this study contained TGFβ1 and/or Nodal growth factors in addition to bFGF to maintain the pluripotent state. There, it is plausible to reason that the observed effects on *NANOG* expression are due to the presence of these growth factors in culture medium. The combination of bFGF and TGFβ growth factors in defined PSC media may enhance the self-renewal capacity and robustness of pluripotency through regulation of *NANOG* expression, compared to the sole use of bFGF (e.g., in undefined PSC culture conditions). It is currently unknown whether the undefined PSC culture media and matrices contained or secreted factors that can promote TGFβ signaling and whether LN-521 can enhance PSC proliferation and self-renewal specifically through *NANOG*. Gene expression analysis indicated that the pluripotency marker *SOX2* was downregulated under defined PSC culture conditions. Since SOX2 is also a neural stem-cell progenitor marker^16^, our observation is in line with the decreased population of differentiating PSCs in the defined PSC culture conditions group. Indeed, when iPSCs cultured under the defined conditions were induced towards differentiation, the protein expression levels of the pluripotency markers POU5F1 and NANOG significantly decreased, whereas that of SOX2 increased. These findings suggested that increased SOX2 expression may serve as an early indicator of differentiation.

We also observed downregulation of well-known pluripotency signaling pathways in defined PSC culture conditions, such as Hippo signaling and the canonical Wnt signaling pathway. The TGFβ signaling pathway was also downregulated under the defined PSC culture conditions. Upon further investigation, it was discovered that the reduced expression of BMP ligands and receptors played a role in this downregulation. No significant differences were observed downstream of TGFβ, Activin and Nodal signaling between the defined and undefined PSC culture conditions. The presence of TGFβ growth factors in the defined rather than undefined PSC medium suggests that the undefined components in PSC culture media, including the culture matrix, may contain and secrete a variety of growth factors that can activate multiple signaling pathways involved in pluripotency and differentiation. However, the number of different components in defined PSC media is more limited, resulting in a reduced utilization of signaling pathways upstream of the POU5F1-NANOG-SOX2 pluripotency network. This restriction in signaling pathway activation likely contributes to the increased homogeneity observed between PSC lines under defined culture conditions, their reproducibility, and the standardization of their differentiation potential.

We demonstrated that the transfer and adaptation of undefined PSCs to defined culture conditions largely reduced the inter-sample heterogeneity. These adapted PSCs showed gene expression patterns that closely resembled those of PSCs originally cultivated under defined culture conditions. This suggests that undefined PSCs can adapt their gene expression patterns upon changing the culture matrix and media, and/or that there is positive selection for undefined PSC populations that utilize similar pluripotency signaling pathways as those in defined PSC culture conditions. Importantly, these results highlight that simply adapting established undefined PSC cultures to defined conditions can greatly reduce the overall variability between PSC lines, and potentially improve their overall quality and experimental reproducibility.

### Calcium signaling and pluripotency

Ca^2+^-binding proteins were significantly overrepresented among the upregulated proteins under defined PSC culture conditions, which led us to investigate Ca^2+^ signaling in iPSCs. Our results demonstrated significant alterations in the spontaneous Ca^2+^ activity of iPSCs during early differentiation, characterized by reduced frequency of Ca^2+^ activity bouts, decreased Ca^2+^ peak amplitude, and increased Ca^2+^ peak rise time. High levels of Ca^2+^ activity have been shown to be specific to and dependent on culture conditions^17^. Elevating intracellular Ca^2+^, either through GPCR agonists or SERCA pump inhibitors, can substitute bFGF in feeder-free human PSC cultures^17^. Thus, high Ca^2+^ activity is required for pluripotency.

Consistent with the observed changes in Ca^2+^ activity during early differentiation, the expression of specific Ca^2+^ binding proteins also showed corresponding changes. Upregulated Ca^2+^-binding proteins in defined PSCs became downregulated during differentiation, and *vice versa*. This suggests potential specificity and relationship between the type of Ca^2+^ activity a PSC displays, the expression of specific Ca^2+^-binding proteins, and the fate of the cells. Similar observations have been made on mouse ESCs, which were shown to display distinct patterns of altered spontaneous Ca^2+^ activity during the ESC-to-neural progenitor and the neural progenitor-to-neuron/astrocyte transition phases^15^. Exit from naïve mESC pluripotency is inhibited by elevated intracellular Ca^2+^ levels^15^. Chelation of cytosolic Ca^2+^ in mESCs induces differentiation by disrupting the expression of *c-Myc*, a transcription factor involved in maintaining PSC self-renewal and pluripotency^18^. Interestingly, *c-Myc* was significantly upregulated in our defined PSC culture condition array dataset, and its activity has been shown to be dependent on direct interaction with the Ca^2+^-binding protein calmodulin^19^.

Thus, unique components of the defined PSC culture conditions may contribute to the generation of spontaneous Ca^2+^ oscillations with highly specific frequencies, widths, spatial distributions, and amplitudes. The functional outcomes of these Ca^2+^ oscillations depend on the binding kinetics of Ca^2+^ to specific effectors (e.g., Ca^2+^ ion-binding proteins such as calmodulin) and by the downstream interactions of these effectors with their unique downstream targets (e.g., transcription factors such as c-Myc) to regulate PSC pluripotency and self-renewal capacity^20^. Smaller decoding of PSC-specific Ca^2+^ oscillations could result in a change in effector activation and downstream repression of pluripotency transcription factors, whereas transcription factors involved in early differentiation may become activated.

### Ca^2+^-binding proteins and the maintenance of pluripotency in defined PSC culture conditions

High expression levels of Ca^2+^-binding proteins in the dataset used here were specifically associated with the pluripotent state. However, it is still unclear whether these proteins actively drive pluripotency or if the type of Ca^2+^ activity displayed by PSCs determines the expression of specific Ca^2+^-binding proteins. The upregulated Ca^2+^-binding proteins identified under the defined PSC culture conditions serve different functions but have not yet been directly linked to the maintenance of PSC pluripotency. Nevertheless, their functions suggest that they may be involved in PSC-dependent Ca^2+^ signaling, self-renewal, and pluripotency. For instance, CALML4 is an essential component of the adhesion complex composed of cadherins and myosin, which are involved in mechanical signal transduction^21^. We demonstrated that blocking cell adhesion disrupts the typical high Ca^2+^ activity displayed by iPSCs under defined culture conditions. During early iPSC differentiation, adhesion-based Ca^2+^ activity is significantly modified, suggesting differential regulation of these mechanisms between differentiating iPSCs and their pluripotent counterparts. AIF1 has been suggested to promote cellular proliferation via the MAPK signaling pathway^22^. Pluripotent cells are characterized by a high proliferation rate that promotes self-renewal. Importantly, the MAPK signaling pathway has been specifically linked to defined PSC culture conditions^23^. Other upregulated Ca^2+^-binding proteins such as PVALB and VSNL1 appear to play a more direct role in Ca^2+^ signaling. PVALB buffers Ca^2+^ ions and its expression has been linked to the maintenance of rapid Ca^2+^ dynamics^24^. Worth noting, the downregulation of PVALB was associated with increased cell body size, which was also observed during early PSC differentiation. VSNL1 is a Ca^2+^ sensing protein involved in signal transduction. It has been shown to promote cell proliferation through the regulation of P2X3/P2Y2 receptors, but also to interact with the α-subunit of nicotinic acetylcholine receptors (nAChRs) and their translocation to the cell surface membrane upon depolarization stimuli^25, 26^. Interestingly, we found an increased population of iPSCs in the pluripotent state that responded to depolarization stimuli compared to those that entered differentiation. This implies that VSNL1-dependent translocation and cell-surface expression of nAChRs upon depolarization stimuli/Ca^2+^ influx may be intact in PSCs cultured under defined conditions.

In conclusion, defined culture conditions led to enhanced inter-PSC sample homogeneity while suppressing differentiation through regulation of Ca^2+^ signalling. A deeper insight into these processes may facilitate the standardization and improvement of defined PSC culture conditions, and thereby clinical applications of human PSCs.

## Acknowledgments

We thank the donors, patients and families. Further, we would like to thank employees of the iPS core facility, Falk and Uhlén laboratory at Karolinska Institutet for support. We thank the BEA facility for support with the Illumina arrays. This study was financed by grants to A.F., VR (2019-01498), hjärnfonden (FO2019-0246, FO2021-0234), VINNOVA (IndiCell 2021-02695), cancerfonden (20 1159 Pj), and to P.U. (VR 2017-00815 and 2021-03108), Hjärnfonden (FO2018-0209 and FO2020-0199), Cancerfonden (19 0544 Pj, 19 0545 Us, and 22 2454 Pj), Barncancerfonden (PR2020-0124 and PR2022-0111). I.E. was supported by a MSCA EF Seal Of Excellence postdoctoral fellowship from the Strategy Group of EU Coordination in Sweden VINNOVA (2021-01834).

## Author contributions

Conceptualization, I.E., M.K., B.U., P.U. and A.F.; methodology, I.E., M.K., M.S., B.U. and D.W.; investigation, I.E., M.K. and B.U., formal analysis, I.E. and B.U.; visualization: I.E. and B.U.; writing – original draft, I.E.; writing – review and editing, I.E., M.K., M.S., B.U., P.U. and A.F.; resources, I.E., B.U., P.U. and A.F.; funding acquisition, I.E., P.U. and A.F.; supervision, P.U. and A.F.

## Inclusion and diversity statement

We support inclusive, diverse, and equitable conduct of research.

## STAR Methods

### Ethical statements

This study was approved by the regional ethical review board in Stockholm, Sweden (Dnr 2016/430-31 and dnr 2012/208-31). Written informed consent was obtained from all donors involved or from their legal guardians.

### Fibroblast culture and reprogramming

Human fibroblast lines were established via mechanical dissection and enzymatic digestion (with dispase and collagenase type IA) of skin biopsies previously obtained from healthy and diagnosed individuals. Fibroblasts were cultured in IMDM (Life technologies), 10% Fetal bovine serum (Invitrogen), 1% NEAA (Gibco) and 1% Penicillin-streptomycin (Life Technologies) on 0.1% gelatin coated plates. To generate iPSCs, approximately 100,000 fibroblasts were transduced using non-integrative Sendai virus vectors encoding the four Yamanaka factors POUF51, SOX2, KLF4 and cMYC (CytoTune iPS Sendai Reprogramming kit, Life Technology) with a multiplicity of infection (MOI) of 3.

### PSC culture conditions

Human iPSCs and ESCs were cultured under either undefined culture conditions or defined and xenofree culture conditions as followed:

#### Undefined culture conditions

After one week, transduced cells were re-plated to irradiated human foreskin fibroblasts (hff, ATCC, CRL-2429) plate (1.5×10^6^ hff on a 100 mm 0.1% gelatin coated dish). PSC-like iPSC colonies became visible after 10 days. Colonies were picked manually using a flame-pulled pasteur pipette and individual iPSC colony clumps were transferred to an irradiated human feeder cell coated 4-well plate 3 weeks post-transduction. To completely clear out the Sendai virus, iPSCs were split manually until passage 10. iPSC lines were cultured in standard human ES medium: Knockout DMEM (Gibco), PEST 1:100, NEAA 1:100, 20% Knockout serum replacement (Gibco) with 8ng ng/ml bFGF (R&D). Medium was changed every day and iPSC colonies were split manually every five to seven days. Extra colonies were frozen in a stem cell bank.

For feeder-free, undefined PSC culture conditions, iPSC lines were manually cut, and pieces were transferred to a matrigel (BD) coated plates in mTeSR Medium (Stemcell Technologies). To clear the culture from hff, iPSC lines were cultured for at least 2 more passages before samples were collected for RNA extraction.

#### Defined culture conditions

For defined feeder-free xeno-free iPSC culture conditions, Sendai virus transduced cells were re-plated on a human recombinant laminin-521 (Biolamina) coated plate (5μg/mL) one week post transduction. Emerging PSC-like colonies were picked manually 3 weeks post transduction, and individual iPSC colony clumps were transferred to a fresh Laminin-521 coated plate in Essential 8™ (E8) medium (ThermoFisher Scientific). From the next and onward passages iPSC lines were split enzymatically using TrypLE Select (ThermoFisher Scientific). Single cells were plated on laminin-521 coated plates in the present of 10 μM Rock inhibitor (Y27632, Millipore). iPSC colonies were plated with the density of 15,000-20,000 cells/cm^2^, split every four to five days, culture medium was replaced daily, and extra cells were frozen in stem cell banks.

### Karyotyping

Karyotyping of iPSCs was done as described in^10^.

### Illumina Gene Expression Array

mRNA was extracted using the RNeasy kit (Qiagen) according to manufacturers protocol. RNA quality and quantity was determined on the Bioanalyzer (Agilent) as in^27^. RNA was amplified according to manufacturer’s protocol using the Illumina TotalPrep RNA Amplification Kit (Ambion). Next, samples were hybridized to an Illumina Gene Expression HT12 Direct Hybridization assay according to manufacturer’s protocol and run on an Illumina BeadChip reader (Illumina HT 12v.4). All raw microarray data is available upon request to the corresponding authors.

### PluriTest

To characterize the pluripotency of the iPSC lines, the microarray iScan idat files were uploaded onto the online bioinformatics tool PluriTest^28^.

Pluritest compares transcriptional profiles of a sample to an extensive transcriptional profile reference set of 223 human embryonic, 41 human iPSCs, somatic cells and tissues. This analysis provides a pluripotency and novelty score. The pluripotency score indicates how strong a model-based pluripotency signature is expressed in the samples analyzed and the novelty score indicates the general model fit for a sample. All iPSCs included in this study passed both scores and were thus determined to be pluripotent (**table S1**).

### Microarray data analysis

Microarray idat files were imported into R^29^ with the miodin package version 0.5.3^30^, using probe annotation file HumanHT-12 version 4.0. The raw microarray data was first normalized with quantile normalization, and technical replicates of each biological sample were then combined by taking the mean across the technical replicates. Differential gene expression analysis was performed with limma^31^ and adjusted for sex, twin and disease status of the samples. P-values were adjusted for multiple testing with Benjamini-Hochberg correction. Genes with absolute log2 fold change > 1 and adjusted p-value < 0.05 were considered significant. Annotation enrichment analysis of biological processes and KEGG pathways was performed with Enrichr^32^ using significant genes from the differential expression analysis as input.

### Protein-protein interaction network analysis

Protein-protein interactions (PPI) between DEGs were obtained via the GeneMANIA plugin in Cytoscape (v.3.9.1) and included physical interactions, predicted interactions, shared protein domains, co-localization and pathways^6^. All interactions were combined and assembled in a reference network and duplicates were removed.

The physical interaction enrichment (PIE) algorithm was used to account for biases in the number of reported protein interactions for disease-associated genes in the generated reference PPI network^7^. PIE scores and associated p-values were calculated against 10 000 random protein groups obtained by number-matched subsamplings selected from the reference PPI network for all DEGs, and as well separately from downregulated DEGs and upregulated DEGs.

### RNA extraction, cDNA synthesis and quantitative PCR

To confirm the expression of DEGs in pluripotent culture conditions, iPSCs cultured for 1 week in either E6 or E8 conditions were selected for reverse transcriptase quantitative (q)PCR. Total RNA was purified from confluent 6-well plates of 5 different iPSC control lines and qPCR were performed as described with the following adaptations^33^. After cell lysis, the samples were purified on a QIAshredder column (Qiagen). Moreover, to avoid genomic DNA contamination, RNA was subjected to DNase treatment using the DNA-free kit (Ambion) before cDNA synthesis. 1μg RNA/sample was reverse transcribed into cDNA using the iScript cDNA (Bio-Rad) according to manufacturers’ procedures.

Quantitative PCRs (qPCRs) were performed using the Fast SYBR Green Master Mix (Life Technologies) on the QuantStudio™ 5 System (Applied Biosystems). The following cycling conditions were used: initial denaturation for 20s at 95°C, followed by 1 s at 95°C and 20 s at 60°C for 40 cycles (QPCR data collection). The products were then denatured at 95°C for 1 s and cooled to 60°C for 20 s (melt curve data collection).

For each cell line, three biological and two technical replicates were analyzed. Differential gene expression was calculated using the 2^ΔΔCt^ method. The average Ct value for each sample was calculated and subtracted from the geometric mean Ct value of the reference genes *GAPDH* and *TBP* to calculate the ΔCt value. All primer sequences for the gene transcripts that were tested in this manuscript are provided in the STAR methods table.

### Flow Cytometry

iPSCs grown in either E6^TM^ or E8^TM^ were washed with PBS, followed by chemical harvesting using TrypLE Select, 1.2 million cells were fixed in 250μl fixation/permeabilization buffer for 15 min at RT using the FOXP3 Transcription Factor Staining Buffer Set (eBioscience, Invitrogen). Cells were washed with permeabilization buffer and stained for 20 min in the dark with the conjugated pluripotency antibodies (1μl/100 000 cells) listed in the STAR methods table. After washing, cells were resuspended in stain buffer and passed through a 35 μm cell strainer (Falcon). All samples were run on a CytoFLEX (Beckman Coulter) and analyzed with FlowJo™ v10.8 Software (BD Life Sciences).

### Calcium imaging

iPSCs grown on a LN-521 substrate in a 35mm culture dish, were loaded with the Ca^2+^-sensitive fluorescence indicator Fluo-4/AM (5 µM; Invitrogen, USA) and Pluronic F-127 (0.625% ThermoFisher Scientific) and incubated for 20 min at 37°C in Krebs-Ringer buffer, containing: NaCl (119 mM), KCl (2.5 mM), NaH_2_PO_4_ (1 mM), CaCl_2_ (2.5 mM), MgCl_2_ (1.3 mM), HEPES (20 mM) and D-Glucose (11 mM), with pH adjusted to 7.4, (All from Sigma). Ca^2+^-measurements were performed in Krebs-Ringer buffer at RT on an upright microscope (Carl Zeiss) equipped with a 20x 1.0 NA lens (Carl Zeiss). Excitation was assessed at 480 nm with a wavelength switcher (DG4, Sutter Instrument) at a sampling frequency of 0.5 Hz. The equipment was controlled with, and data was collected using MetaFluor (Molecular Devices). FIJI, MATLAB (R2021a, MathWorks, USA) and FluoroSNNAP^34^ were used to process and analyze the collected data.

### Statistical analysis

The results were plotted in R or PRISM (GraphPad Prism 9). Statistical analysis of live Ca^2+^ imaging and qPCR was done with a non-parametric Kruskal-Wallis test in combination with a Dunns multiple comparisons post-test. Non-parametric Mann-Whitney test was used to calculate differences between two conditions. P-values lower than 0.05 were considered as significant, with * p < 0.05, ** p < 0.01, *** p < 0.001, **** p < 0.0001.

## Supplemental information

The supplemental information includes four figures and one table.

## References

1. Takahashi, K., and Yamanaka, S. (2006). Induction of pluripotent stem cells from mouse embryonic and adult fibroblast cultures by defined factors. Cell 126, 663–676. 10.1016/j.cell.2006.07.024.

2. Uhlin, E., Marin Navarro, A., Ronnholm, H., Day, K., Kele, M., and Falk, A. (2017). Integration Free Derivation of Human Induced Pluripotent Stem Cells Using Laminin 521 Matrix. J Vis Exp. 10.3791/56146.

3. Albalushi, H., Kurek, M., Karlsson, L., Landreh, L., Kjartansdottir, K.R., Soder, O., Hovatta, O., and Stukenborg, J.B. (2018). Laminin 521 Stabilizes the Pluripotency Expression Pattern of Human Embryonic Stem Cells Initially Derived on Feeder Cells. Stem Cells Int 2018, 7127042. 10.1155/2018/7127042.

4. Dutan Polit, L., Eidhof I., McNeill R.V., Warre-Cornish K.M., Ohki C.M.Y., Monet Walter N., Sala C., Verpelli C., Radtke F., Galderisi S., Mucci A., Collo G., Edenhofer F., Castrén M.L., Réthelyi J.M., Ejlersen M., Hohmann S.S., Ilieva M.S., Lukjanska R., Matuleviciute R., Michel T. M., de Vrij F.M.S., Kushner S.A., Lendemeijer B., Kittel-Schneider S., Ziegler G.C., Gruber-Schoffnegger D., Pasterkamp J., Kasri A., Potier M., Knoblich J.A., Brüstle O., Peitz M., Merlo Pich E., Harwood A.J., Abranches E., Falk A., Vernon AC., Edna Grünblatt E., Srivastava D.P. (2023). Recommendations, guidelines, and best practice for the use of human induced pluripotent stem cells for neuropharmacological studies of neuropsychiatric disorders. Neuroscience Applied 2. 10.1016/j.nsa.2023.101125.

5. Uhlin, E., Ronnholm, H., Day, K., Kele, M., Tammimies, K., Bolte, S., and Falk, A. (2017). Derivation of human iPS cell lines from monozygotic twins in defined and xeno free conditions. Stem Cell Res 18, 22–25. 10.1016/j.scr.2016.12.006.

6. Montojo, J., Zuberi, K., Rodriguez, H., Kazi, F., Wright, G., Donaldson, S.L., Morris, Q., and Bader, G.D. (2010). GeneMANIA Cytoscape plugin: fast gene function predictions on the desktop. Bioinformatics 26, 2927–2928. 10.1093/bioinformatics/btq562.

7. Eidhof, I., van de Warrenburg, B.P., and Schenck, A. (2019). Integrative network and brain expression analysis reveals mechanistic modules in ataxia. J Med Genet 56, 283–292. 10.1136/jmedgenet-2018-105703.

8. Wu, S., Johansson, J., Damdimopoulou, P., Shahsavani, M., Falk, A., Hovatta, O., and Rising, A. (2014). Spider silk for xeno-free long-term self-renewal and differentiation of human pluripotent stem cells. Biomaterials 35, 8496–8502. 10.1016/j.biomaterials.2014.06.039.

9. Lam, M., Moslem, M., Bryois, J., Pronk, R.J., Uhlin, E., Ellstrom, I.D., Laan, L., Olive, J., Morse, R., Ronnholm, H., et al. (2019). Single cell analysis of autism patient with bi-allelic NRXN1-alpha deletion reveals skewed fate choice in neural progenitors and impaired neuronal functionality. Exp Cell Res 383, 111469. 10.1016/j.yexcr.2019.06.014.

10. Kele, M., Day, K., Ronnholm, H., Schuster, J., Dahl, N., and Falk, A. (2016). Generation of human iPS cell line CTL07-II from human fibroblasts, under defined and xeno-free conditions. Stem Cell Res 17, 474–478. 10.1016/j.scr.2016.09.028.

11. Plaza Reyes, A., Petrus-Reurer, S., Padrell Sanchez, S., Kumar, P., Douagi, I., Bartuma, H., Aronsson, M., Westman, S., Lardner, E., Andre, H., et al. (2020). Identification of cell surface markers and establishment of monolayer differentiation to retinal pigment epithelial cells. Nat Commun 11, 1609. 10.1038/s41467-020-15326-5.

12. Arjun McKinney, A., Petrova, R., and Panagiotakos, G. (2022). Calcium and activity-dependent signaling in the developing cerebral cortex. Development 149. 10.1242/dev.198853.

13. Ma, M., Hua, S., Min, X., Wang, L., Li, J., Wu, P., Liang, H., Zhang, B., Chen, X., and Xiang, S. (2022). p53 positively regulates the proliferation of hepatic progenitor cells promoted by laminin-521. Signal Transduct Target Ther 7, 290. 10.1038/s41392-022-01107-7.

14. Penton, C.M., Badarinarayana, V., Prisco, J., Powers, E., Pincus, M., Allen, R.E., and August, P.R. (2016). Laminin 521 maintains differentiation potential of mouse and human satellite cell-derived myoblasts during long-term culture expansion. Skelet Muscle 6, 44. 10.1186/s13395-016-0116-4.

15. Liu, B., Chen, S., Xu, Y., Lyu, Y., Wang, J., Du, Y., Sun, Y., Liu, H., Zhou, H., Lai, W., et al. (2021). Chemically defined and xeno-free culture condition for human extended pluripotent stem cells. Nat Commun 12, 3017. 10.1038/s41467-021-23320-8.

16. Xu, R.H., Sampsell-Barron, T.L., Gu, F., Root, S., Peck, R.M., Pan, G., Yu, J., Antosiewicz-Bourget, J., Tian, S., Stewart, R., and Thomson, J.A. (2008). NANOG is a direct target of TGFbeta/activin-mediated SMAD signaling in human ESCs. Cell Stem Cell 3, 196–206. 10.1016/j.stem.2008.07.001.

17. Apati, A., Berecz, T., and Sarkadi, B. (2016). Calcium signaling in human pluripotent stem cells. Cell Calcium 59, 117–123. 10.1016/j.ceca.2016.01.005.

18. Ermakov, A., Daks, A., Fedorova, O., Shuvalov, O., and Barlev, N.A. (2018). Ca(2+) - depended signaling pathways regulate self-renewal and pluripotency of stem cells. Cell Biol Int 42, 1086–1096. 10.1002/cbin.10998.

19. Raffeiner, P., Schraffl, A., Schwarz, T., Rock, R., Ledolter, K., Hartl, M., Konrat, R., Stefan, E., and Bister, K. (2017). Calcium-dependent binding of Myc to calmodulin. Oncotarget 8, 3327–3343. 10.18632/oncotarget.13759.

20. Snoeck, H.W. (2020). Calcium regulation of stem cells. EMBO Rep 21, e50028. 10.15252/embr.202050028.

21. Kapustina, M., and Cheney, R.E. (2020). A new light chain for myosin-7. J Biol Chem 295, 9297–9298. 10.1074/jbc.H120.014595.

22. Li, Q., Hu, L., Liu, G., Yin, X., Li, Y., Wei, X., Duan, N., Zhao, X., Gong, Q., and Du, Z. (2023). Inhibition of AIF-1 alleviates laser-induced macular neovascularization by inhibiting endothelial cell proliferation via restrained p44/42 MAPK signaling pathway. Exp Eye Res 231, 109474. 10.1016/j.exer.2023.109474.

23. Nakashima, Y., and Omasa, T. (2016). What Kind of Signaling Maintains Pluripotency and Viability in Human-Induced Pluripotent Stem Cells Cultured on Laminin-511 with Serum-Free Medium? Biores Open Access 5, 84–93. 10.1089/biores.2016.0001.

24. Klocke, B., Krone, K., Tornes, J., Moore, C., Ott, H., and Pitychoutis, P.M. (2023). Insights into the role of intracellular calcium signaling in the neurobiology of neurodevelopmental disorders. Front Neurosci 17, 1093099. 10.3389/fnins.2023.1093099.

25. Li, C., Pan, W., Braunewell, K.H., and Ames, J.B. (2011). Structural analysis of Mg2+ and Ca2+ binding, myristoylation, and dimerization of the neuronal calcium sensor and visinin-like protein 1 (VILIP-1). J Biol Chem 286, 6354–6366. 10.1074/jbc.M110.173724.

26. Williams, T.A., Monticone, S., Crudo, V., Warth, R., Veglio, F., and Mulatero, P. (2012). Visinin-like 1 is upregulated in aldosterone-producing adenomas with KCNJ5 mutations and protects from calcium-induced apoptosis. Hypertension 59, 833–839. 10.1161/HYPERTENSIONAHA.111.188532.

27. Shahsavani, M., Pronk, R.J., Falk, R., Lam, M., Moslem, M., Linker, S.B., Salma, J., Day, K., Schuster, J., Anderlid, B.M., et al. (2018). An in vitro model of lissencephaly: expanding the role of DCX during neurogenesis. Mol Psychiatry 23, 1674–1684. 10.1038/mp.2017.175.

28. Muller, F.J., Schuldt, B.M., Williams, R., Mason, D., Altun, G., Papapetrou, E.P., Danner, S., Goldmann, J.E., Herbst, A., Schmidt, N.O., et al. (2011). A bioinformatic assay for pluripotency in human cells. Nat Methods 8, 315–317. 10.1038/nmeth.1580.

29. R Core Team (2021). R: A Language and Environment for Statistical Computing. R foundation for Statistical computing, Vienna, Austria.

30. Ulfenborg, B. (2019). Vertical and horizontal integration of multi-omics data with miodin. BMC Bioinformatics 20, 649. 10.1186/s12859-019-3224-4.

31. Ritchie, M.E., Phipson, B., Wu, D., Hu, Y., Law, C.W., Shi, W., and Smyth, G.K. (2015). limma powers differential expression analyses for RNA-sequencing and microarray studies. Nucleic Acids Res 43, e47. 10.1093/nar/gkv007.

32. Kuleshov, M.V., Jones, M.R., Rouillard, A.D., Fernandez, N.F., Duan, Q., Wang, Z., Koplev, S., Jenkins, S.L., Jagodnik, K.M., Lachmann, A., et al. (2016). Enrichr: a comprehensive gene set enrichment analysis web server 2016 update. Nucleic Acids Res 44, W90–97. 10.1093/nar/gkw377.

33. Eidhof, I., Baets, J., Kamsteeg, E.J., Deconinck, T., van Ninhuijs, L., Martin, J.J., Schule, R., Zuchner, S., De Jonghe, P., Schenck, A., and van de Warrenburg, B.P. (2018). GDAP2 mutations implicate susceptibility to cellular stress in a new form of cerebellar ataxia. Brain 141, 2592–2604. 10.1093/brain/awy198.

34. Patel, T.P., Man, K., Firestein, B.L., and Meaney, D.F. (2015). Automated quantification of neuronal networks and single-cell calcium dynamics using calcium imaging. J Neurosci Methods 243, 26–38. 10.1016/j.jneumeth.2015.01.020.

