## Supplementary information for "Defined human PSC culture conditions robustly maintain human PSC pluripotency through Ca^2+^ signaling"

### Author list footnotes

[+]present address: BioLamina AB, Sundbyberg, 172 66, Sweden

[#]present address: Department of Molecular Medicine and Surgery and Center for Molecular Medicine, Karolinska Institutet, Stockholm, 171 11, Sweden

### Supplemental information

**Supplementary table 1: An overview of and detailed information on the PSC samples included in the analysis.**

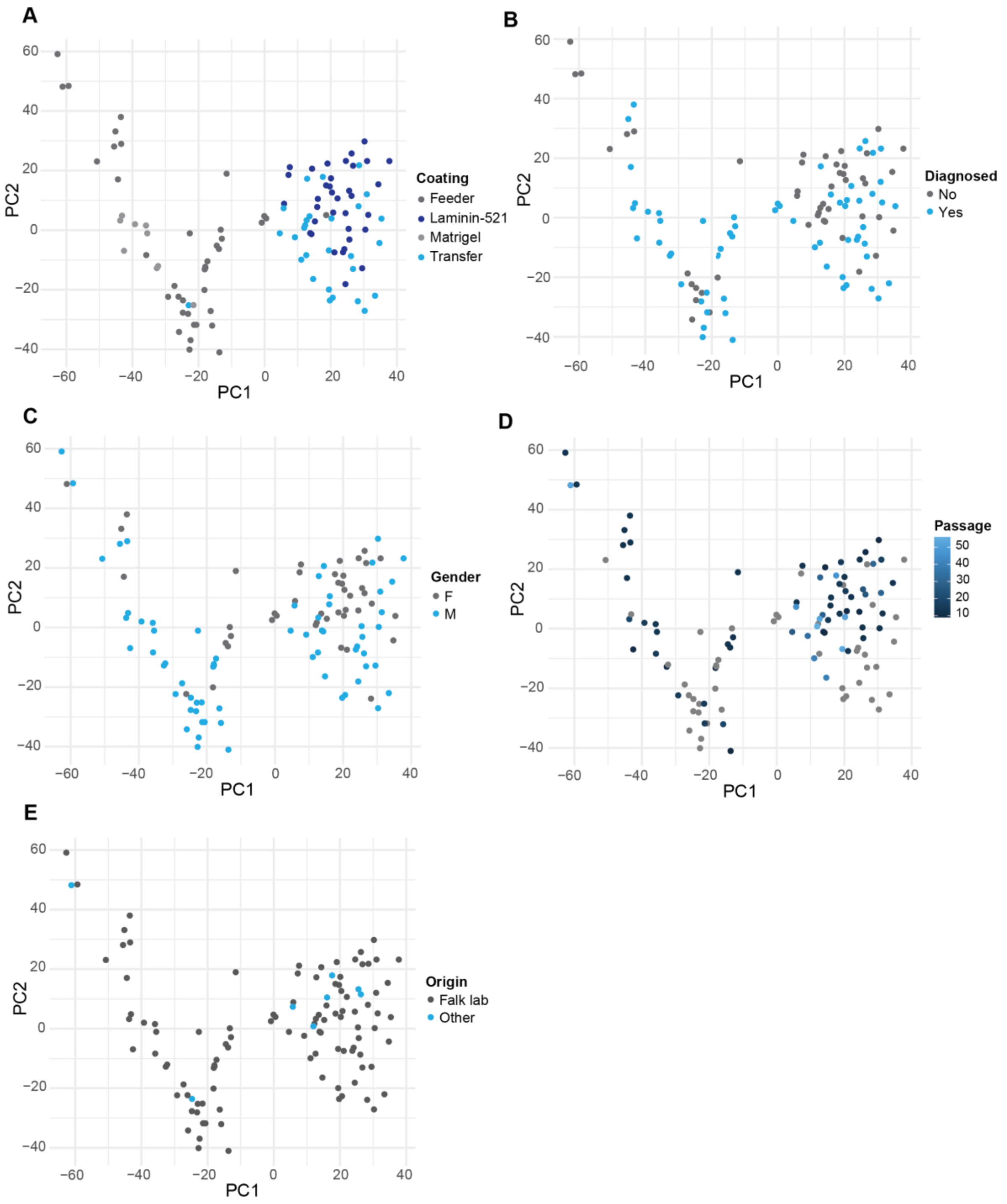


**Figure S1: PCA clustering is unaffected by potential confounding variables in the dataset. A.** Visualization of plate coating in the PCA plot: human feeders (in dark grey), Laminin-521 (in dark blue), Matrigel (in light grey) and Transfer from feeders to Laminin-521 (in light blue). **B.** Visualization of iPSCs that originate from individuals with (in blue) or without (in grey) a diagnosis in the PCA plot. **C.** Visualization of PSCs that originate from male (in blue) or female (in grey) individuals in the PCA plot. **D.** Visualization of the passage number of the PSC line at the time of the microarray in the PCA plot. **E.** Visualization of the laboratory origin of the PSC line in the PCA plot.

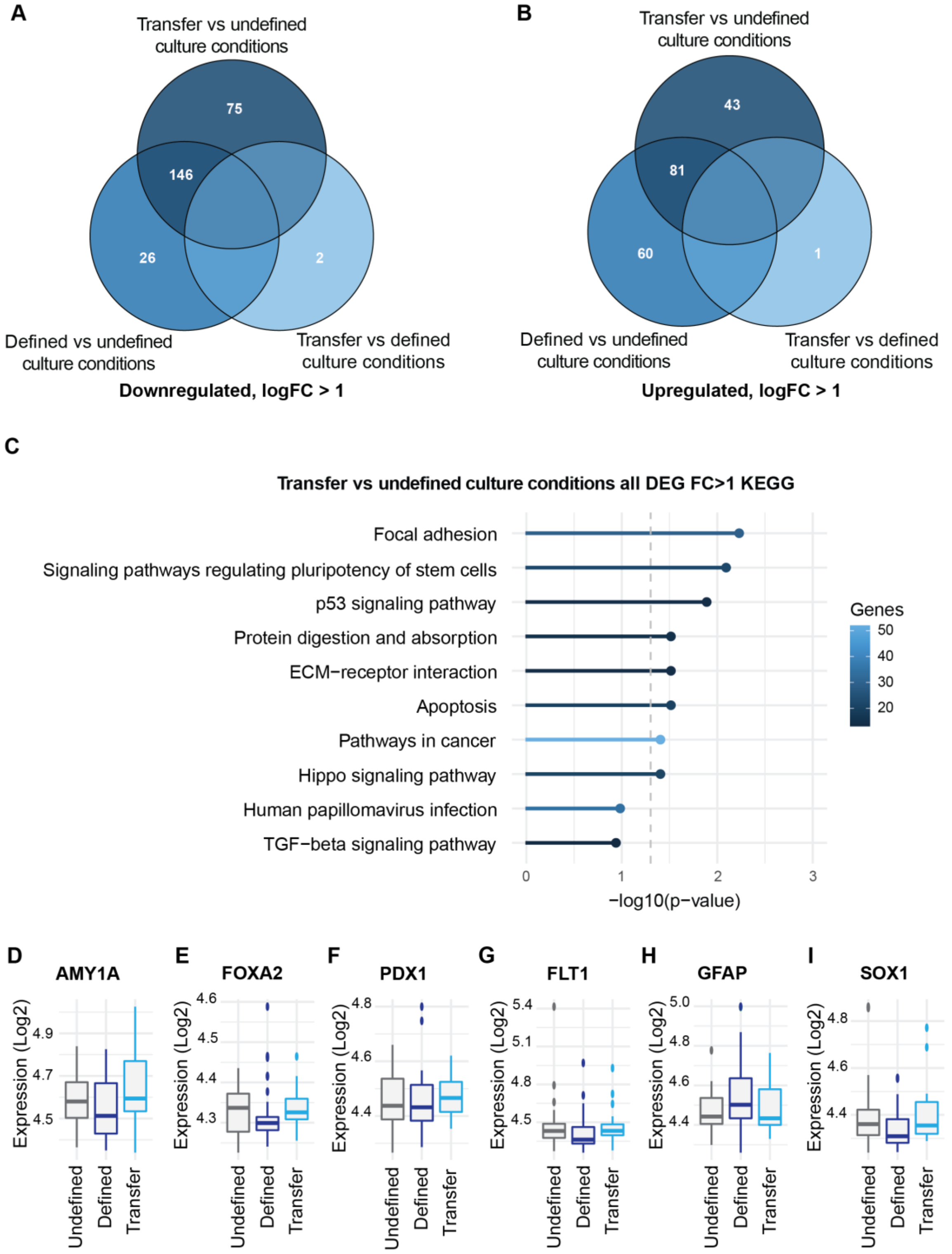


**Figure S2. A.** VENN diagram showing the total number of significantly downregulated genes passing a log foldchange > 1 and between the indicated PSC culture conditions. **B.** VENN diagram showing the total number of significantly upregulated genes passing a log foldchange > 1 between the indicated PSC culture conditions. **C.** KEGG pathway analysis on DEGs between PSCs established in undefined culture conditions that were transferred to defined culture conditions and undefined PSC culture conditions. **D-I.** Log2 microarray expression values of the endodermal markers *AMY1A* (**D**), *FOXA2* (**E**) and *PDX1* (F), mesodermal marker *FLT1* (**G**), and ectodermal markers *GFAP* (**H**) and *SOX1* (**I**) for the indicated PSC culture conditions.

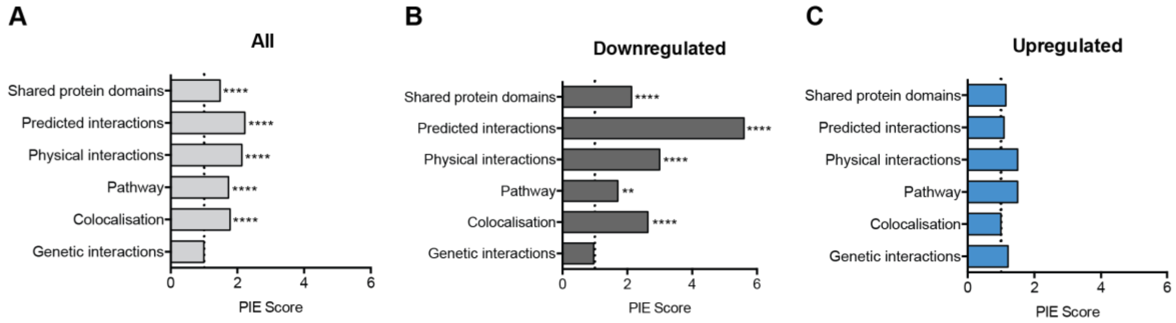


**Figure S3:** Physical Interaction Enrichment Score analysis performed separately for the indicated protein-protein interaction data for all DEGs (**A**), downregulated DEGs (**B**) and upregulated DEGs (**C**) between defined and undefined PSC culture conditions. **p < 0.01, ****p < 0.0001, based on 10,000 repetitions.

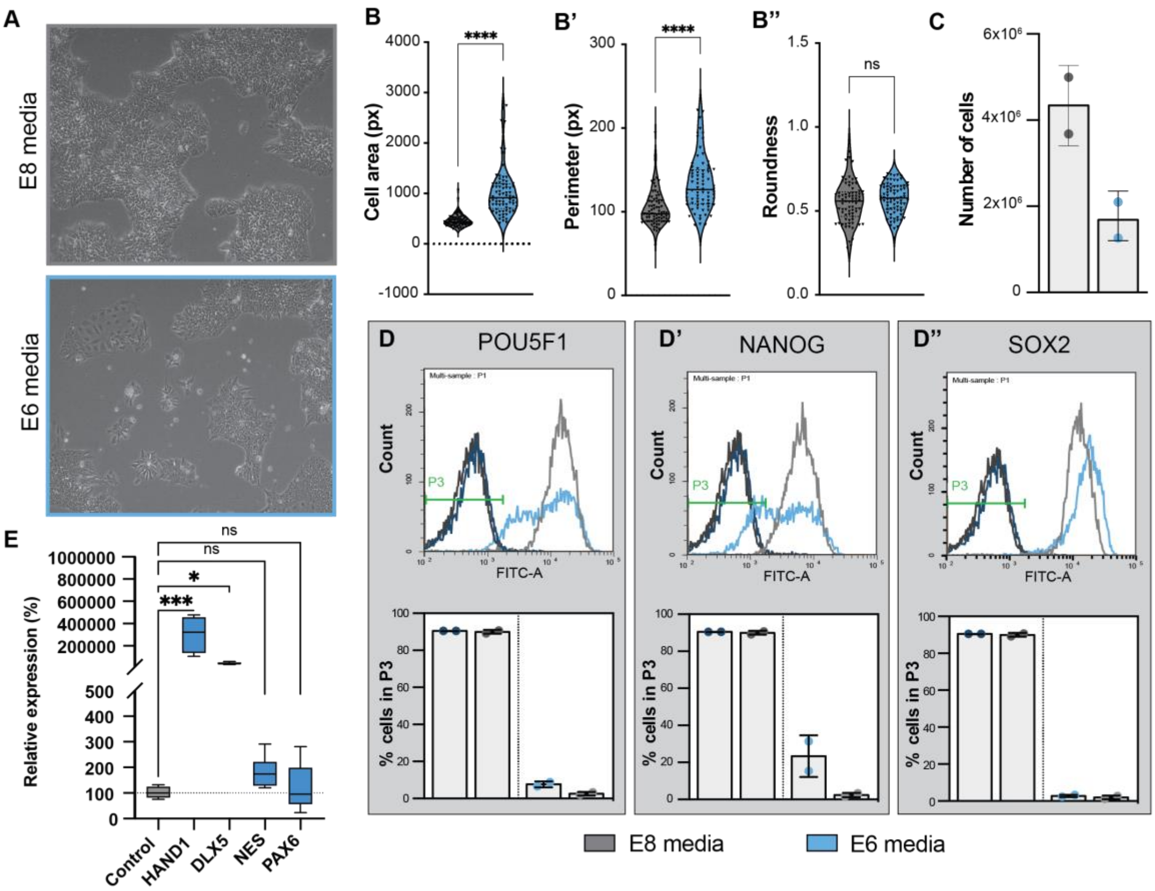


**Figure S4: Culturing iPSCs for 7 days in defined E6 culture media, without TGFβ and bFGF, induces differentiation and mimic undefined culture conditions. A.** Bright field images of iPSC colonies cultured in defined E8 media (grey) and E6 media (blue), lacking TGFβ and bFGF. **B.** FIJI quantification of iPSC morphology parameters in E8 versus E6 media. B: iPSC area (in pixel), B' iPSC perimeter (in pixel). B'' Roundness. **C.** Quantification of the number of iPSCs after 1 week in E8 (in grey) or E6 (in blue) culture media. **D.** FACS analysis of transcription factors POU5F1 (D), NANOG (D') and SOX2 (D'') that make up the core pluripotency network. Light blue: iPSCs cultured in E6 media stained for the indicated marker. The bargraphs below show the quantification of the percentage of cells that lay in the area of IgM stained cells (Green area P3 in D). Dark grey: iPSCs cultured in E8 stained with negative control: IgM. Dark blue: iPSCs cultured in E6 stained with negative control: IgM. Light grey: iPSCs cultured in E8 media stained for the indicated marker. **E.** Relative gene expression data (%) of ectodermal and mesodermal transcription factors in E6 media (data is normalized against the E8 control condition). Pooled data of 5 different iPSC control lines, line at median. *$p < 0.05$, ***$p < 0.001$, ****$p < 0.0001$, ns: not significant.
